# A novel reporter to visualize and quantify endogenous inflammasomes and caspase-1 recruitment

**DOI:** 10.1101/2025.03.30.646205

**Authors:** Florian N. Gohr, Maria H. Christensen, Lea-Marie Jenster, Yonas M. Tesfamariam, Dorothee J. Lapp, Jason Mackenzie, Florian I. Schmidt

## Abstract

Inflammasomes are signaling complexes that coordinate inflammation by inducing rapid cytokine secretion and that protect against invading pathogens by initiating death of infected cells. Yet, current methods do not allow us to specifically monitor inflammasome activation in real-time at endogenous protein levels. To overcome this shortcoming, we developed and characterized a novel fluorescent reporter that visualizes inflammasome assembly and reports on the recruitment of the effector protein caspase-1 without impairing downstream signaling. The reporter permits the analysis of inflammasome assembly over time as well as precise quantification of cells with assembled inflammasomes. We have successfully applied the reporter in lentivirus-transduced human and murine cell lines and primary cells but have also applied recombinant viruses that encode the reporter in their genome to detect inflammasome responses in primary cells. Mouse intestinal enteroids expressing the reporter allowed us to visualize how Salmonella infection triggers NAIP/NLRC4 inflammasome assembly, cell death, and expulsion in complex tissues. Lastly, we apply the new tool to gain mechanistic insights and prove that caspase-1 activation on inflammasomes relies on the assembly of filaments of caspase-1^CARD^, which can be terminated by the CARD-only protein CARD17 to shut down cytokine secretion. We anticipate broad application of the reporter in fundamental and applied research, as it permits the quantitative assessment of inflammasome assembly at both high temporal and spatial resolution or in high throughput.

## Introduction

Inflammasomes are large macromolecular signaling complexes that coordinate the inflammation, which is needed to clear infections, but contributes to disease if not controlled appropriately (Martinon, Burns and Tschopp, 2002; Broz and Dixit, 2016). Despite the central role of inflammasomes in immunity, important aspects of inflammasome biology remain elusive. In particular, we do not sufficiently understand how inflammasome activation in a tissue is coordinated with other signaling pathways, which cell types can assemble inflammasomes, and what governs the timing and regulation of inflammasome assembly at the single cell level. A fundamental bottleneck for our current understanding of inflammasomes is the lack of tools to reliably quantify inflammasome assembly under physiological conditions. The ideal readout for inflammasome activation would work 1) at the single cell level, 2) at endogenous levels of all inflammasome components, 3) in living cells, 4) in (human) primary cells, and 5) as upstream as possible to avoid pathogen-mediated countermeasures to inflammasome signaling. Yet, most currently applied techniques to quantify inflammasome assembly rely on bulk measurements of cytokines or lactate dehydrogenase (LDH) activity in the supernatant as well as caspase-1-mediated protein processing in cell lysates.

In distinct sentinel cell types, inflammasomes are nucleated by a handful of different sensors, which oligomerize in response to cytosolic insults (Broz and Dixit, 2016). All inflammasome sensors either contain a caspase recruitment domain (CARD) or a pyrin domain (PYD). They engage the adaptor protein ASC through homotypic CARD:CARD or PYD:PYD interactions, which nucleates the polymerization of ASC^PYD^ filaments, which are cross-linked by CARD:CARD interactions to yield spheric structures that can be visualized as ASC specks by immunofluorescence staining with ASC antibodies (Stutz *et al*., 2013; Florian I Schmidt *et al*., 2016). Unprocessed caspase-1 is comprised of a CARD, and the large (p20) and small (p10) catalytic subunits. Caspase-1 is recruited to ASC specks through ASC^CARD^:caspase-1^CARD^ interactions, where two molecules of unprocessed caspase-1 (p46) dimerize through the catalytic subunits (Ross *et al*., 2022; Schmidt, 2023). Caspase-1 p46 subsequently undergoes autoproteolytic cleavage of the interdomain linker (IDL) between the catalytic subunits, yielding dimers composed of two p33 (CARD + large catalytic subunit) and two p10 (small catalytic subunit) fragments. The resulting tetrameric (p33/p10)_2_ complex appears to have maximal catalytic activity. Initial activation is followed by cleavage of the CARD linker (CDL) between the CARD and the catalytic domains, resulting in the release of (p20/p10)_2_ from ASC specks and the gradual loss of enzymatic activity (Boucher *et al*., 2018). As enzymatically active caspase-1 involves the homo-dimerization of catalytic subunits, caspase-1 activation is widely described to rely on caspase-1 dimerization. In the simplest model, dimerization is merely facilitated by the proximity of many caspase-1 molecules on ASC specks (Ross *et al*., 2022). *In vitro*, however, seeds of recombinantly produced ASC^CARD^ nucleate the polymerization of caspase-1^CARD^ into highly ordered filaments (Lu *et al*., 2014), implying that this highly cooperative process governs caspase-1 activation. While the structure of such CARD filaments was determined by cryo-electron microscopy (Lu *et al*., 2016), it has not been experimentally addressed whether this also occurs in cells. Evidently, the mechanism of caspase-1 activation has important implications: Caspase-1 activation by dimerization on ASC specks might be limited by a finite number of binding sites in cells with low ASC expression, while activation along caspase-1^CARD^ filaments would allow for continuous recruitment, independent of the initial number of ASC molecules.

As we currently lack the tools necessary to visualize and quantify inflammasome assembly in suitable experimental conditions, we developed a new, non-invasive inflammasome reporter with low background that is compatible with microscopy, flow cytometry, and cell sorting. Moreover, it directly reports on the recruitment of caspase-1, which allows us to investigate the mode of caspase-1 recruitment and enables the visualization of ASC-independent caspase-1 recruitment. We apply the reporter to prove that caspase-1 activation involves the assembly of caspase-1 filaments in cells, which offers an additional layer of regulation by the CARD-only protein CARD17.

## Results

### Caspase-1^CARD^-EGFP is efficiently recruited to ASC specks with negligible background

Canonical inflammasomes are characterized by the assembly of ASC into a large complex, which becomes apparent as an ‘ASC speck’ when stained with ASC antibodies. While this reports on the endogenous signalosome, this approach requires fixation, permeabilization, and staining. This precludes live cell experiments and suffers from the fragility of pyroptotic cells. A commonly used genetically encoded reporter is the ectopic expression of ASC-EGFP (Figure 1A (Stutz *et al*., 2013)), which changes the endogenous levels of ASC or—in the most extreme case—renders cell types lacking ASC competent for inflammasome assembly. Importantly, overexpressed ASC can oligomerize in the absence of upstream sensor activation, requiring careful selection of cell lines and expression levels. As currently available techniques do not allow the easy visualization and monitoring of inflammasome responses, we developed a novel reporter that fulfills the following criteria: 1) it does not require staining; 2) it identifies inflammasome activation in living or fixed cells; 3) it is suitable for primary cells and tissues; 4) it offers single cell resolution; 5) it lends itself to quantification by microscopy and flow cytometry; 6) it exhibits low background; 7) and it reports on the recruitment of caspase-1 as the most relevant outcome of ASC speck assembly.

**Figure 1:**
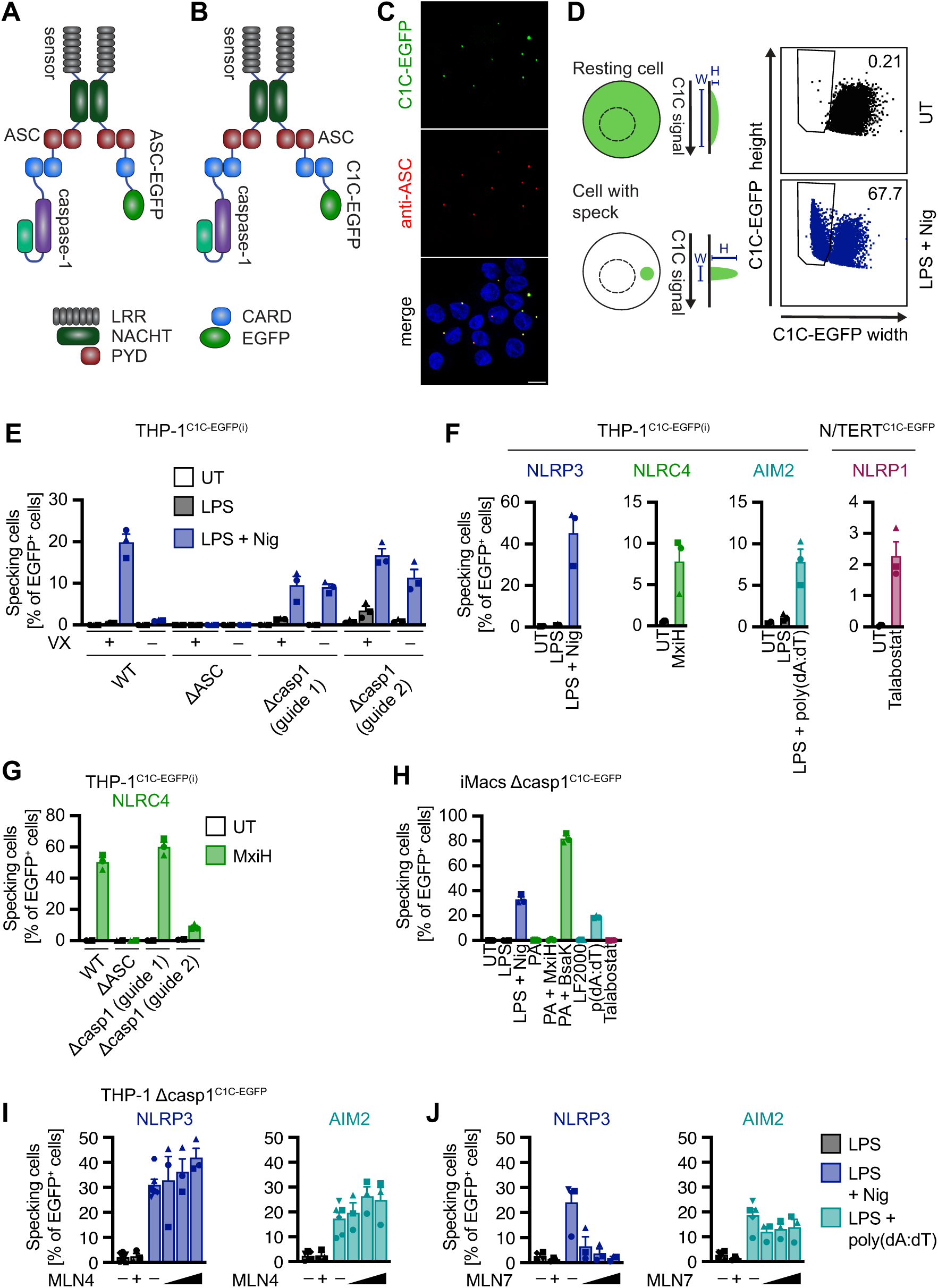
C1C-EGFP allows detection of inflammasome assembly. **A–B)** Schematic representation of the inflammasome reporters ASC-EGFP (A) and caspase-1^CARD^(C1C)-EGFP (B). ASC-EGFP is incorporated into nascent ASC specks via PYD:PYD or CARD:CARD interactions with the sensor and endogenous ASC. C1C-EGFP is recruited to nascent ASC specks via CARD:CARD interactions with ASC or endogenous caspase-1. **C)** PMA-differentiated THP-1^C1C-EGFP(i)^ cells were induced with 1 µg/mL doxycycline (dox) for 24 h and treated with 200 ng/mL LPS for 3 h and 10 µM nigericin (Nig) for 1 h. 40 µM VX-765 was added 15 min before and during nigericin treatment. Cells were fixed, stained for ASC and DNA, and cells were recorded by confocal microscopy. One representative image is shown. Scale bar: 10 μm. **D)** THP-1^C1C-EGFP^ cells were differentiated with PMA and treated with LPS and nigericin as in C. UT: untreated. Cells with ASC specks recruiting C1C-EGFP from a representative experiment were identified by flow cytometry using the altered width and height signal of C1C-EGFP as illustrated. **E)** THP-1^C1C-EGFP(i)^, THP-1 ΔASC^C1C-EGFP(i)^ or THP-1 Δcaspase-1(casp1)^C1C-EGFP(i)^ cells were differentiated with PMA and treated with 200 ng/mL LPS for 3 h and 10 µM nigericin for 1 h in the presence or absence of 40 µM VX-765. The fraction of specking cells was measured by flow cytometry. **F)** PMA-differentiated THP-1^C1C-EGFP(i)^ cells or N/TERT-1^C1C-EGFP^ keratinocytes were treated as indicated in the presence of 40 µM (THP-1, short stimulation) or 100 µM (N/TERT-1, long stimulation) VX-765. The fraction of specking cells was measured by flow cytometry. NLRP3 was activated with 200 ng/mL LPS for 3 h and 10 µM nigericin for 1 h; NLRC4 was stimulated with 0.1 µg/mL LFn-MxiH and 1 µg/mL PA for 1 h; for AIM2 activation, cells were pre-treated with LPS for 3 h, transfected with 1 µg/mL poly(dA:dT) using Lipofectamine^TM^ 2000, and incubated for 4 h; NLRP1 was stimulated with 30 µM talabostat for 20 h. The fraction of specking cells was measured by flow cytometry. **G)** THP-1^C1C-EGFP(i)^, THP-1 ΔASC^C1C-EGFP(i)^ or THP-1 Δcasp1^C1C-EGFP(i)^ cells were differentiated with PMA and treated with LFn-MxiH and PA as described in F. The fraction of specking cells was measured by flow cytometry. **H**) iMac caspase-1^−/−mC1C-EGFP^ cells were treated with LPS and nigericin, PA and LFn-MxiH or 0.1 µg/mL LFn-BsaK, poly(dA:dT), or talabostat as described in F. The fraction of specking cells was measured by flow cytometry. **I–J)** THP-1 Δcasp^C1C-EGFP(i)^ cells were stimulated with LPS and nigericin (NLRP3) or poly(dA:dT) (AIM2) in the presence of 0, 5, 10, or 20 µM MLN4924 (MLN4, I) or MLN7243 (MLN7, J). Cells treated with LPS alone were cultivated in the absence (-) or presence (+) of 20 µM MLN4 or MLN7, respectively. All data represents mean values from 3 independent experiments ± SEM.

As the downstream effects of inflammasomes depend on the recruitment of caspase-1 via its CARD, we wondered whether caspase-1^CARD^ (C1C) could be used to recapitulate caspase-1 recruitment. We genetically fused C1C to EGFP, expecting that C1C-EGFP would be recruited to the inflammasome akin to full-length caspase-1 (Figure 1B). To verify that C1C-EGFP can identify ASC specks, we generated a THP-1 cell line that doxycycline-inducibly expresses C1C-EGFP (THP-1^C1C-EGFP(i)^). We activated NLRP3 by treatment with LPS and nigericin, stained for endogenous ASC, and compared the distribution of both molecules by microscopy (Figure 1C). C1C-EGFP redistributed into specks and completely colocalized with endogenous ASC, thus confirming that the reporter visualizes *bona fide* inflammasomes. While it is possible to quantify ASC speck assembly and C1C-EGFP recruitment by fluorescence microscopy and image analysis (see below), we were also keen to measure inflammasome formation by flow cytometry, as a faster and more scalable way of quantification suitable for larger cell and sample numbers. Sester *et al*. had pioneered a flow cytometry-based method for inflammasome detection (Sester *et al*., 2015), which relies on the relocalization of ASC during inflammasome formation: compared to the diffusely distributed ASC signal in resting cells, the signal of ASC specks in activated cells has a smaller diameter, but higher intensity. In flow cytometry, this change translates into a decreased fluorescent pulse width along with an increased signal height (Figure 1D). Since microscopy revealed a redistribution of C1C-EGFP equivalent to that of ASC, we tested whether this assay works equally well with C1C-EGFP instead of ASC. We measured LPS- and nigericin-treated THP-1^C1C-EGFP(i)^ cells by flow cytometry and plotted the GFP height against the GFP width. Indeed, we observed a well separated population of cells exhibiting the characteristic behavior of cells with ASC specks. Importantly, we found that THP-1 cells with assembled inflammasomes could only be revealed by flow cytometry if caspase-1 was inhibited by VX-765 treatment, or when experiments were conducted in caspase-1 knockout (KO) cells (Figure 1E). For this reason, all subsequent experiments were conducted in the presence of VX-765 unless otherwise stated. We assume that caspase-1-catalyzed cleavage of GSDMD and the ensuing pyroptosis renders cells too fragile to sustain trypsinization, pipetting steps, and sedimentation. In line with that, cells that cannot form GSDMD pores due to the expression of antagonistic GSDMD nanobodies can be analyzed by flow cytometry in the absence of caspase-1 inhibitors (Schiffelers *et al*., 2024).

C1C-EGFP is typically only recruited to existing ASC specks as it cannot bind most inflammasome sensors directly. We therefore expected that its recruitment, and thus the readout, is independent of the upstream sensor. To test this, we stimulated THP-1^C1C-EGFP(i)^ for activation of different inflammasome sensors (Figure 1F). NLRP3 was activated by priming with LPS, followed by stimulation with nigericin. To activate NAIP/NLRC4, we delivered MxiH (*Shigella flexneri* needle protein) into the cytosol using the anthrax toxin delivery system: MxiH was fused to the anthrax lethal factor N-terminus (LFn) and administered together with the *Bacillus anthracis* protective antigen (PA) (Milne *et al*., 1995). AIM2 was activated by transfecting poly(dA:dT). To test NLRP1 inflammasome assembly, we generated N/TERT-1 keratinocytes expressing C1C-EGFP (N/TERT-1^C1C-EGFP^). We treated these cells with talabostat (also known as Val-boroPro). All reporter cells showed robust specking in response to the respective triggers, as quantified by flow cytometry (Figures 1D, F). As expected, no C1C-EGFP specks were detected after NLRP3 activation in the absence of ASC, but recruitment of C1C-EGFP to ASC specks was observed in the absence of endogenous caspase-1 (Figure 1E). NLRC4 can directly recruit caspase-1^CARD^ *in vitro* (Zhang *et al*., 2015) and mouse NLRC4 has been shown to induce pyroptosis in the absence of ASC (Broz *et al*., 2010). Yet, we did not observe the formation of specks—or other structures—in THP-1 cells lacking ASC (Figure 1G). This observation suggests that human NLRC4 depends on ASC for recruiting and activating caspase-1. Accordingly, we have previously shown that NLRC4 activation did not lead to LDH release in ΔASC THP-1 cells (Schiffelers *et al*., 2024).

To make the reporter available for research in mice, we generated mouse caspase-1^CARD^ (mC1C)-EGFP. We introduced this reporter into iMacs, an immortalized mouse macrophage cells line from caspase-1 KO animals (iMacs casp-1^−/−mC1C-EGFP(i)^), and stimulated NLRP3, NAIP1/NLRC4, and AIM2 (Figure 1H). All treatments resulted in a clear specking response, confirming the functionality of the reporter system in murine cells. These data show that C1C-EGFP is a robust reporter that provides single cell resolution for inflammasome assembly in microscopy and flow cytometry without the need for staining.

ASC-EGFP is prone to undergo PYD and CARD-mediated polymerization to assemble specks even in the absence of activated inflammasome sensors. After extensive use of our C1C-based reporters, we never encountered issues with background in any cell line. To compare the background levels of both reporters in a relevant cell line expressing endogenous ASC, THP-1 cells were transduced with lentivirus encoding ASC-EGFP or C1C-EGFP under the control of different constitutive promoters with varying strengths. Transduced cells were differentiated with PMA and rested for two days in the presence of the caspase-1 inhibitor VX-765 to avoid the loss of specking cells due to pyroptosis. All cells were then harvested, and both the reporter expression and the fraction of specking cells were measured and evaluated (Figure S1). High expression of ASC-EGFP under the control of the CMV promoter (Figure S1A) resulted in a high background of ASC specks in the absence of inflammasome triggers (Figure S1B). The background in all other samples was negligible. This experiment shows that C1C-EGFP is less prone to generate background specking in the absence of genuine inflammasome activators and thus is compatible with a wide range of promoters and expression system.

### C1C-EGFP permits easy screening of inflammasome inhibitors

Drugs that target inflammasome assembly are highly desirable. IL-1β-based readouts integrate a multitude of effects besides inflammasome assembly, including expression levels of caspase-1 and IL-1β, caspase-1 activity, and GSDMD pore formation. This impedes the search for inhibitors that specifically target inflammasomes as well as accurate interpretation of results. A more proximal readout would therefore be beneficial. To test C1C-EGFP as a tool to screen for potential inflammasome inhibitors, we treated THP-1 Δcasp-1^C1C-EGFP(i)^ cells with triggers for NLRP3 (nigericin) or AIM2 (poly(dA:dT)), (Figures 1I–J) in the presence of increasing concentrations of different inhibitors of the ubiquitination machinery.

MLN4924 (Pevonedistat) inhibits NEDD8-activating enzyme and, thereby, ubiquitination by Cullin-RING ubiquitin ligases. We previously found that NLRP1 activation depends on this kind of ubiquitin ligases (L. M. Jenster *et al*., 2023). However, NLRP3 and AIM2 were not inhibited be MLN4924 (Figure 1I). MLN7243 (TAK-243) inhibits the ubiquitin activating enzyme E1. It completely abrogated NLRP3 inflammasome formation at 20 μM, while AIM2 was not affected, indicating that NLRP3 but not AIM2 activation depends on ubiquitination (Figure 1J). Taken together, these proof-of-concept experiments show that the C1C reporter is suitable to screen for drugs inhibiting inflammasome assembly for fundamental research or therapy.

### C1C-EGFP does not interfere with inflammasome signaling

C1C-EGFP is recruited to inflammasomes using the same interactions as caspase-1 itself, which raises the question whether it competes with endogenous caspase-1 for recruitment and thereby hampers its recruitment and activation. To assess this, we compared WT THP-1 macrophages with THP-1^C1C-EGFP^ and THP-1^C1C-EGFP(i)^. Expression of the reporter C1C-EGFP was comparable under the constitutive ubiquitin C promoter (pUbC) in THP-1^C1C-EGFP^ cells and after doxycycline induction in THP-1^C1C-EGFP(i)^ (Figure S2A). We next determined LPS-mediated activation of TLR4 by measuring tumor necrosis factor (TNF) secretion, and observed that cytokine secretion, and thus LPS priming, was not impaired in any cell line (Figure S2B). Cells expressing C1C-EGFP, whether induced or constitutive, assembled specks after treatment with nigericin or MxiH (Figure 2A). The inflammasome response to nigericin was stronger after LPS priming as expected. The amount of released IL-1β into the medium was comparable to WT THP-1 cells in all cell lines and therefore not reduced by the expression of C1C-EGFP (Figure 2B). When we compared NLRP3 activation with LPS and nigericin to NLRC4 activation with MxiH, we found that both the specking response and IL-1β release were stronger after NLRP3 activation, indicating that both readouts were consistent. Interestingly, treatment of unprimed and LPS-primed THP-1 cells with nigericin yielded robust C1C specking responses in both conditions, while IL-1β release was substantially lower in the absence of priming, likely due to limited pro-IL-1β expression in the absence of priming. This highlights an advantage of assessing inflammasomes on the basis of assembly, as it is independent of the upregulation of pro-IL-1β.

**Figure 2:**
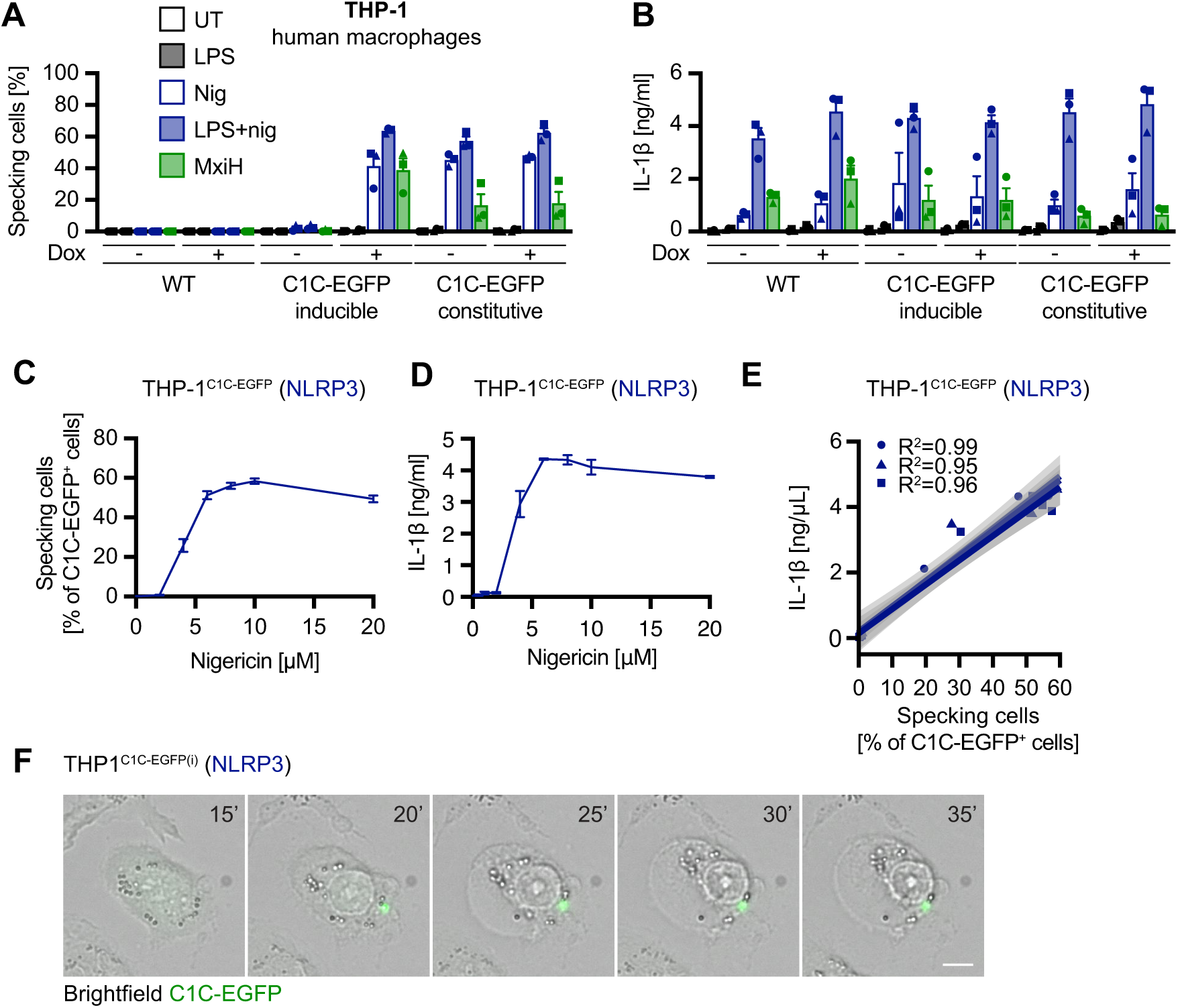
C1C-EGFP does not interfere with inflammasome signaling. **A**–**B)** PMA-differentiated THP-1, THP-1^C1C-EGFP^, or THP-1^C1C-EGFP(i)^ were treated with or without doxycycline as indicated and stimulated with LPS and nigericin or with PA + LFn-MxiH for 1 h as described in Fig. 1F. Cells for flow cytometry (A) were stimulated in the presence of 40 µM VX-765. The fraction of specking cells was quantified by flow cytometry (A) and IL-1β release was quantified by HTRF (B). **C**–**E)** PMA-differentiated THP-1^C1C-EGFP^ were treated with 200 ng/mL LPS for 3 h and the indicated concentrations of nigericin for 1 h. Cells for flow cytometry (C) were stimulated in presence of 40 µM VX-765. The fraction of specking cells was quantified by flow cytometry (C) and secreted IL-1β quantified by HTRF (D). The linear correlation for 3 independent experiments with confidence band 0.95 is plotted in (E). **F)** PMA-differentiated THP-1^C1C-EGFP(i)^ cells were treated with 200 ng/mL LPS for 3 h and 10 µM nigericin. Live cells were recorded using widefield microscopy. The number indicates the time point in minutes. Scale bar: 50 µm. Data represents mean values from 3 independent experiments ± SEM.

To assess the correlation between specking frequency and IL-1β release, we treated THP-1^C1C-EGFP^ cells with LPS and different concentrations of nigericin to compare dose-dependent responses (Figures 2C-E). Clear inflammasome activation was observed starting with 4 μM nigericin and plateaued at around 10 μM (our standard working concentration). A linear regression between the fraction of specking cells and IL-1β resulted in a good correlation with R2 of well over 90 % (Figure 2E). This result indicates that the amount of cytokine secreted is primarily dependent on the number of cells assembling inflammasomes. This is in line with the observed binary response to inflammasome triggers in macrophages: Cooperative assembly of inflammasomes either leads to the full response, including cell death and cytokine release, or no response at all. Importantly, this distinction cannot be made on the basis of bulk measurements of cell death or cytokine release.

Ultimately, the formation of inflammasomes leads to pyroptosis, which is associated with swelling of the affected cells due to the influx of water through GSDMD pores. Using live-cell imaging, we also show that specking cells undergo pyroptosis in the presence of C1C-EGFP (Figure 2F). This also highlights the possibility to use the reporter in living cells to investigate the dynamics of inflammasome signaling.

In summary, we show that the detection of ASC specks via C1C-EGFP is a robust method to detect inflammasome signaling that does not interfere with the pathway and is consistent with well-established methods like cytokine measurement.

### C1C-EGFP permits visualization of inflammasomes in primary cells

Primary macrophages are of particular interest in inflammasome research. They can be cultivated long enough to allow genetic manipulation and thus reporter expression. We differentiated primary human monocytes into macrophages using M-CSF and introduced C1C-EGFP into the cells by lentiviral transduction (Figure 3A). To overcome macrophage restriction of the incoming lentivirus by SAMHD1, we produced the lentivirus in cells expressing a fusion of the accessory proteins Vpx from simian immunodeficiency virus (SIV) and Vpr from human immunodeficiency virus 1 (HIV-1) (Sunseri *et al*., 2011; Nodder and Gummuluru, 2019). While Vpr mediates packaging into virions, Vpx induces the degradation of SAMHD in target cells. We next treated the transduced cells with MxiH and assessed C1C-EGFP expression as well as speck formation by flow cytometry. 60% of the cells expressed C1C-EGFP (Figure 3B). We observed that around 80% of C1C-EGFP-positive cells assembled C1C-EGFP specks in response to MxiH, illustrating that C1C-EGFP can be used to detect inflammasomes in genetically modified primary cells.

**Figure 3:**
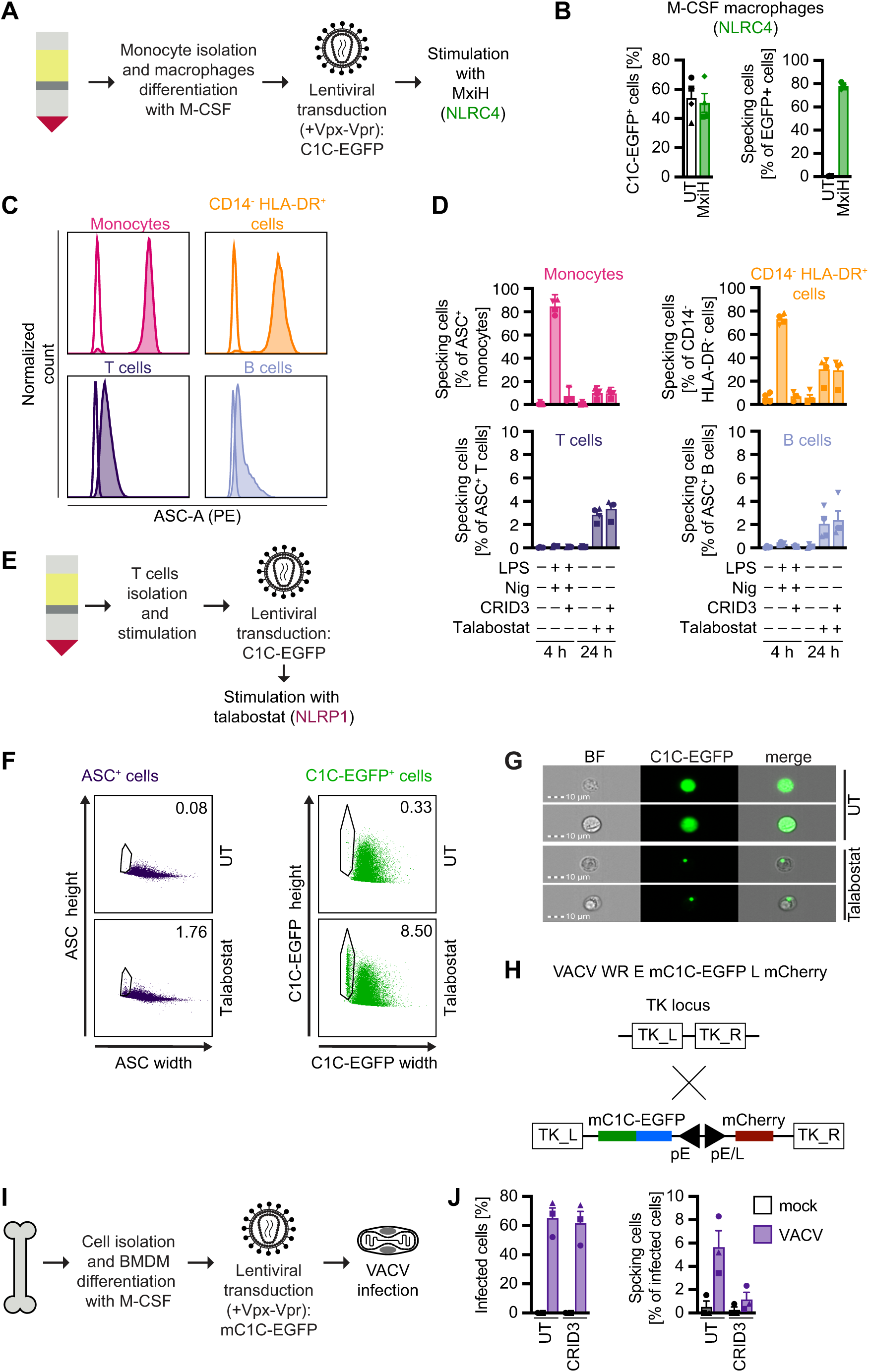
C1C-EGFP reveals inflammasome responses in primary cells. **A**–**B)** Inflammasome responses in human monocyte-derived M-CSF macrophages revealed with C1C-EGFP. Primary human monocytes isolated from buffy coats were differentiated into macrophages with 1 µg/mL M-CSF for 3 days, seeded into 24-well plates, and transduced with 3x concentrated lentivirus packaging Vpx-Vpr and encoding C1C-EGFP (A). After 2 days, the cells were treated with MxiH for 1 h in the presence of VX-765 as described in Fig. 1F. The fraction of C1C-EGFP positive cells and specking cells was quantified by flow cytometry (B). Data represents mean values from 4 donors ± SEM. **C**–**D)** Inflammasome responses in human PBMCs revealed by ASC staining. Human PBMCs were isolated from fresh blood and stimulated in the presence of 100 µM VX-765. Cells were left untreated or treated with either 200 ng/mL LPS for 3 h and 10 μM nigericin for 1 h, or 30 μM talabostat for 24 h. Where indicated, stimulations were done in the presence of 2.5 μM CRID3. Living cells were stained to gate for monocytes (CD14^+^), T cells (CD14^−^ CD3^+^), B cells (CD14^−^ CD19^+^), and CD14^−^ CD3^−^ C19^−^ HLA-DR^+^ cells. Cells were subsequently fixed, permeabilized, additionally stained for ASC, and analyzed by flow cytometry. A representative histogram of ASC staining of unstimulated cells is displayed in (C). The fraction of cells with ASC specks in the indicated subpopulations was quantified by flow cytometry (D). Data represents mean values from 4 donors ± SEM. **E**–**G)** Inflammasome responses in human T cells revealed with C1C-EGFP. CD3^+^ T cells were isolated from fresh blood, stimulated with coated anti-CD3/CD28 antibodies, and transduced with lentivirus encoding C1C-EGFP. After 5 days, transduced T cells were stimulated with 30 μM talabostat for 24 h in the presence of 100 µM VX-765 (E). T cells were stained for ASC to compare ASC speck formation detected by staining of endogenous ASC and C1C-EGFP reporter. ASC speck formation in ASC^+^ or C1C-EGFP^+^ cells was analyzed by flow cytometry (F) or imaging flow cytometry (G). Representative dot plots or images are shown. **H**–**J)** Genome structure of recombinant vaccinia virus (VACV) WR strain expressing murine C1C-EGFP (mC1C-EGFP) from the VACV J2R early promoter and mCherry from a synthetic poxvirus early/late promoter (H). Transgenes were inserted into the TK locus of VACV by homologous recombination. BMDMs from C57BL/6N mice were infected with VACV^E:mC1C-EGFP L:mCherry^ (MOI=25) for 20 h in the presence of 100 µM VX-765, where indicated with 2.5 µM CRID3 (I). Data represents mean values from 3 independent experiments ± SEM.

To investigate which other blood cells might have the potential to form inflammasomes, we stained ASC in peripheral blood mononuclear cells (PBMCs) and found that monocytes and a small population of CD14^−^ HLA-DR^+^ cells expressed high amounts of ASC, while T and B cells expressed ASC to a much lower extent (Figure 3C). To test which cells were able to assemble NLRP1 or NLRP3 inflammasomes, we treated PBMCs with talabostat or LPS and nigericin, respectively. We observed a robust NLRP3 inflammasome response in monocytes and CD14^−^ HLA-DR^+^ cells, but not the other cell types (Figure 3D). Interestingly, in response to talabostat, we observed clear inflammasome assembly in T cells, while responses were more variable in B cells (Figure 3D). To follow up on inflammasome assembly in T cells, we established a protocol to isolate T cells from PBMCs, expand them by anti-CD3/CD28 stimulation, and lentivirally transduce them to express C1C-EGFP (Figure 3E). To validate the inflammasome reporter in T cells, we additionally stained ASC in talabostat-treated cells and compared the specking response as detected by ASC staining and C1C-EGFP (Figure 3F). The low expression of ASC in T cells hampered clear gating on specking cells because the two populations are poorly separated, presumably due to smaller ASC specks than in myeloid cells. The C1C-EGFP signal, however, was substantially stronger than the endogenous ASC signal, offering a better separation between cells with and without specks (Figure 3F). Using image stream analysis, we confirmed the characteristic speck morphology in talabostat-treated C1C-EGFP-expressing T cells (Figure 3G).

Talabostat activates NLRP1, but also CARD8, by displacing DPP9 from autoinhibited complexes of the sensor (Johnson *et al*., 2018; Taabazuing, Griswold and Bachovchin, 2020). To confirm that T cells form NLRP1 inflammasomes, we used a nanobody-based system to directly and specifically activate NLRP1 by ubiquitination that we had described previously (L. M. Jenster *et al*., 2023). In brief, a nanobody (VHH) against the NLRP1^PYD^ was genetically fused to Von Hippel-Lindau factor (VHL), and thus recruits Cullin-2 ubiquitin ligase complexes to ubiquitinate the NLRP1^PYD^, followed by degradation of the NLRP1 N-terminal fragment and the release of the active NLRP1^UPA-CARD^ fragment (Figure S3A). We transduced anti-CD3/anti-CD28-stimulated T cells with lentiviral vectors to simultaneously express C1C-EGFP and VHL-VHH fusions under the control of a bidirectional doxycycline-inducible promoter (Figure S3A). Expression of the nanobody fusion protein directed against NLRP1^PYD^ (VHL-VHH_NLRP1 PYD-1_), but not the control VHH targeting the influenza nucleoprotein (VHL-VHH_NP_), induced a robust specking response in T cells (Figure S3B). The inflammasome response was substantially stronger than after talabostat treatment. In the presence of VX-765, around 20% of the C1C-EGFP-expressing T cells formed specks 8 h post-induction. In the absence of VX-765, T cells expressing C1C-EGFP together with VHL-VHH_PYD-1_ rounded up with the typical morphology of pyroptotic cells after doxycycline induction and could not be detected by flow cytometry, suggesting that the NLRP1-activating T cells indeed die by pyroptosis (Figures S3B–C). Of note, no cytokine secretion by T cells was observed (data not shown).

To study inflammasome responses to virus infections, we decided to generate reporter viruses that themselves encode and express C1C-EGFP in infected cells: this allows quantification of infection and inflammasome assembly by flow cytometry in the same experiment. Moreover, this approach leads to expression of the reporter in cells that cannot easily be genetically manipulated, such as many primary cell types. As a proof of concept, we infected mouse bone marrow-derived macrophages (mBMDM) with a recombinant vaccinia virus (VACV) strain that encodes C1C-EGFP under the control of the VACV J2R early promoter and mCherry under the control of a synthetic poxvirus early/late promoter, both inserted into the TK locus (Figures 3H–I). Approximately 60% of the cells were infected and about 5% of the infected mBMDMs readily formed specks after 6 h (Figure 3J). This response was almost completely abolished in the presence of the NLRP3 inhibitor CRID3 (also known as MCC950), suggesting that inflammasomes were nucleated by NLRP3.

Taken together, these results demonstrate that the new C1C-EGFP reporter works robustly and without any background to detect inflammasomes in primary cells. It could be effectively used to detect inflammasome assembly in primary macrophages and T cells. In addition, it can be encoded in the genomes of recombinant viruses to assess the ability of these viruses to activate an inflammasome response. This approach can be employed in primary cells and potentially lends itself to comparable experiments *in vivo*.

### C1C-EGFP permits visualization of inflammasomes in tissues

Inflammasomes serve important functions in many cell types. While most studies have focused on myeloid cell types, typically in pure cultures, inflammasomes also coordinate inflammation in various other tissues, including epithelia, which are crucial for rapid responses to invading pathogens. Several mouse studies have established a role for NLRC4 inflammasomes in intestinal epithelial cells (IECs). NAIP/NLRC4 in mouse IECs is important in the defense against Salmonella (Sellin *et al*., 2014). The epithelial integrity is preserved by the expulsion of affected cells into the gut lumen (Sellin *et al*., 2014; Rauch *et al*., 2017), which is achieved by myosin-dependent focal contractions within the epithelium (Samperio Ventayol *et al*., 2021).

To test our reporter in intestinal epithelia, we generated mouse intestinal enteroids (MIE) from the small intestine and lentivirally transduced them to express the mC1C-EGFP reporter. To induce inflammasome formation in these cells, we treated the cells with MxiH, a ligand for mouse NAIP1. Additionally, we stained endogenous ASC and visualized the MIEs by confocal microscopy (Figure 4A). The mC1C-EGFP signal colocalized with that of endogenous ASC staining, confirming *bona fide* ASC specks in intestinal epithelia. Next, we tested if inflammasome assembly in MIEs can also be quantified by flow cytometry as described before. To avoid pyroptotic cell death, we treated the MIEs with the caspase-1 inhibitor VX-765 or the pan-caspase inhibitor Z-VAD(OMe)-FMK. After stimulation with MxiH, we dissociated the enteroids into single cells by trypsinization and measured the fluorescence by flow cytometry (Figure 4B). Cells with assembled inflammasome could easily be identified using our standard gating strategy. MxiH induced a robust inflammasome response in ∼40% of enteroid cells. This demonstrates that the NAIP1/NLRC4 inflammasome is active in MIEs and that the reporter is functional.

**Figure 4:**
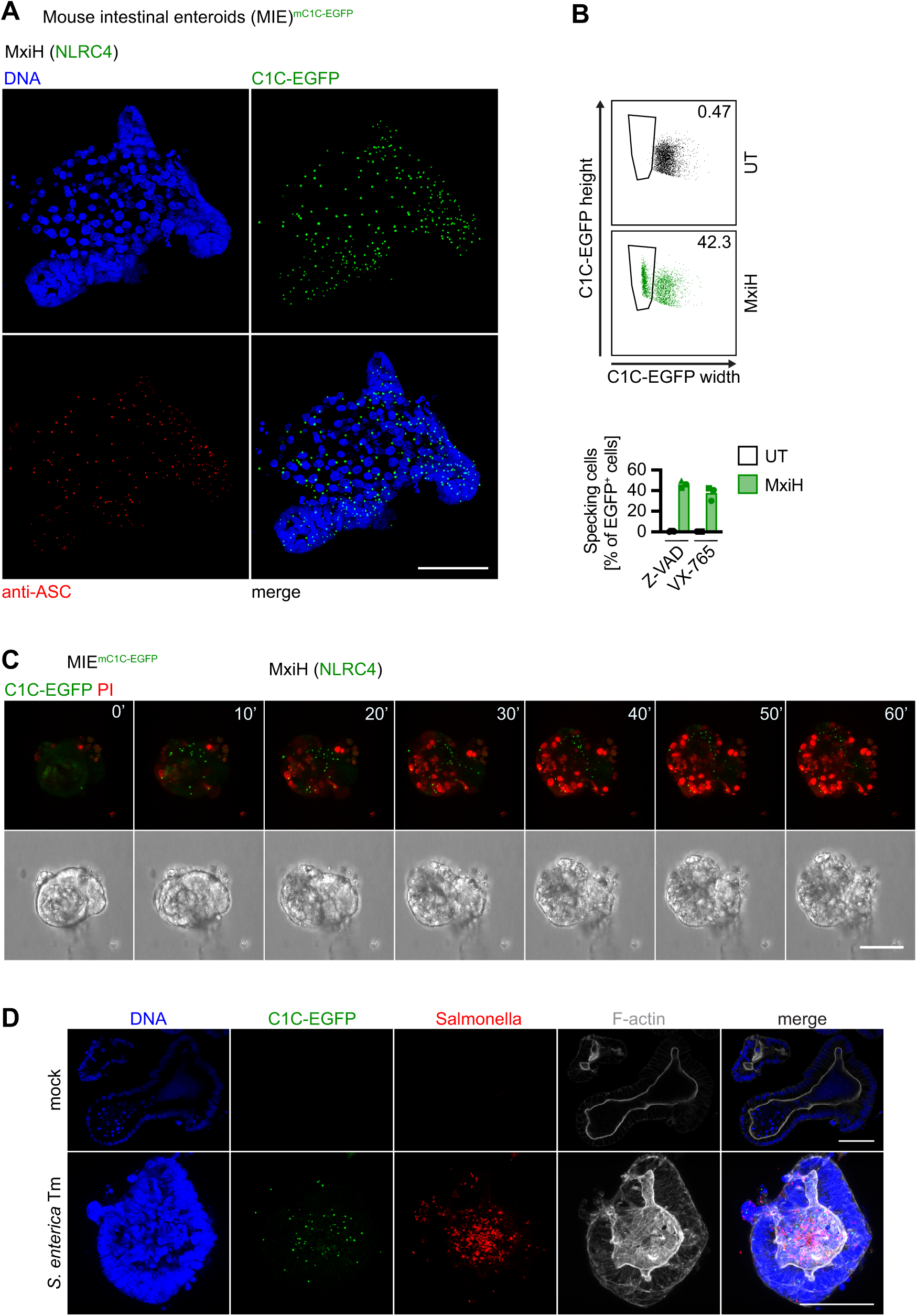
C1C-EGFP permits monitoring of inflammasome assembly and cell extrusion in mouse intestinal enteroids (MIE). **A)** Mouse intestinal enteroids expressing mC1C-EGFP (MIE^mC1C-EGFP^) were treated with 0.1 ug/mL LFn-MxiH and 1 µg/mL PA in the presence of 50 µM Z-VAD(OMe)-FMK for 1 h. Enteroids were fixed, permeabilized, and stained for ASC and DNA. Z-stacks were recorded by confocal microscopy and the maximum intensity projection of a representative enteroid is displayed. The scale bar represents 50 μm. **B)** MIE^mC1C-EGFP^ were treated as in A, but dissociated by trypsinization after stimulation. Single cell suspensions were measured by flow cytometry and a representative dot plot is displayed alongside quantified data from 3 independent experiments ± SEM. **C)** MIE^mC1C-EGF^ were treated with MxiH and PA in DMEM without phenol red and in the presence of 3.33 µg/mL PI as a viability dye. Living cells were recorded by confocal microscopy over time. The maximum intensity projection of a Z-stack of a representative enteroid is shown. Time points are indicated as minutes; the scale bar represents 50 μm. **D)** MIE^mC1C-EGFP^ were infected with *S.* Typhimurium expressing mCherry for 2 h. Cells were fixed and stained for DNA and F-actin. Z projections were recorded by confocal microscopy and the maximum intensity projection of a representative enteroid is shown. The scale bars represent 50 μm.

To capture the dynamics of inflammasome activation in IECs, we used confocal live-cell imaging (Figure 4C). C1C-EGFP specks were already formed within the first 10 minutes after addition of MxiH. One hour after treatment, the enteroids exhibited extensive cell death, as revealed by propidium iodide (PI) uptake and loss of morphological integrity. Interestingly, we could also observe the expulsion of specking cells into the enteroid lumen.

It has previously been reported that mouse IECs form NAIP/NLRC4-dependent inflammasomes in response to infection with enteric bacteria (Sellin *et al*., 2014; Rauch *et al*., 2017). However, the assembly of a *bona fide* inflammasomes has so far not been visualized. To test whether specks are formed in response to bacterial infection, we infected MIE^mC1C-EGFP^ with *Samonella enterica* serovar Typhimurium for 2 h and examined speck formation by confocal microscopy (Figure 4D). We observed robust inflammasome assembly at this early stage of infection with salmonella. Interestingly, most specking cells were found within the lumen of the enteroids, presumably due to expulsion. In contrast to the simultaneous stimulation of NAIP1/NLRC4 in all cells with MxiH, infected enteroids maintained overall integrity over the duration of the experiment, likely because fewer cells were infected and assembled inflammasomes.

These results prove the utility of C1C-EGFP as an inflammasome reporter in a complex three-dimensional cell culture system as well as infection experiments. The results encourage its use in other organoid models and tissues. We confirmed published findings on mouse intestinal NLRC4 inflammasomes and showed the assembly of macromolecular ASC specks as caspase-1 recruitment platforms in IECs.

### C1C-EGFP recapitulates the recruitment and activation of caspase-1 by polymerization in living cells

In contrast to ASC-based reporters, C1C-EGFP not only indicates the assembly of ASC specks, but also recapitulates the recruitment of caspase-1. Recombinantly expressed C1C can assemble into helical filaments *in vitro* using three asymmetric interfaces for CARD:CARD interaction (Lu *et al*., 2016). In such filaments, each C1C subunit interacts with six neighboring subunits using three types of interfaces (type I, II, III, Figure 5A). The end of the filament in which polymerization is nucleated is termed the a-end, while the distal end that is elongated by newly recruited subunits is termed the b-end (Lu *et al*., 2016). Each subunit is recruited via its a-side and recruits further subunits via its b-side. However, it is currently unclear whether activation of caspase-1 on intracellular ASC specks indeed relies on the polymerization of caspase-1^CARD^, as this process has thus far only been observed using recombinant protein. Recent reviews either assume that caspase-1 dimerizes through CARD dimers or proximity (Ross *et al*., 2022) or extrapolate from *in vitro* data that caspase-1 indeed polymerizes through CARD filaments (Figure 5B). C1C-EGFP provides a suitable experimental system to assess this question in intact cells.

**Figure 5:**
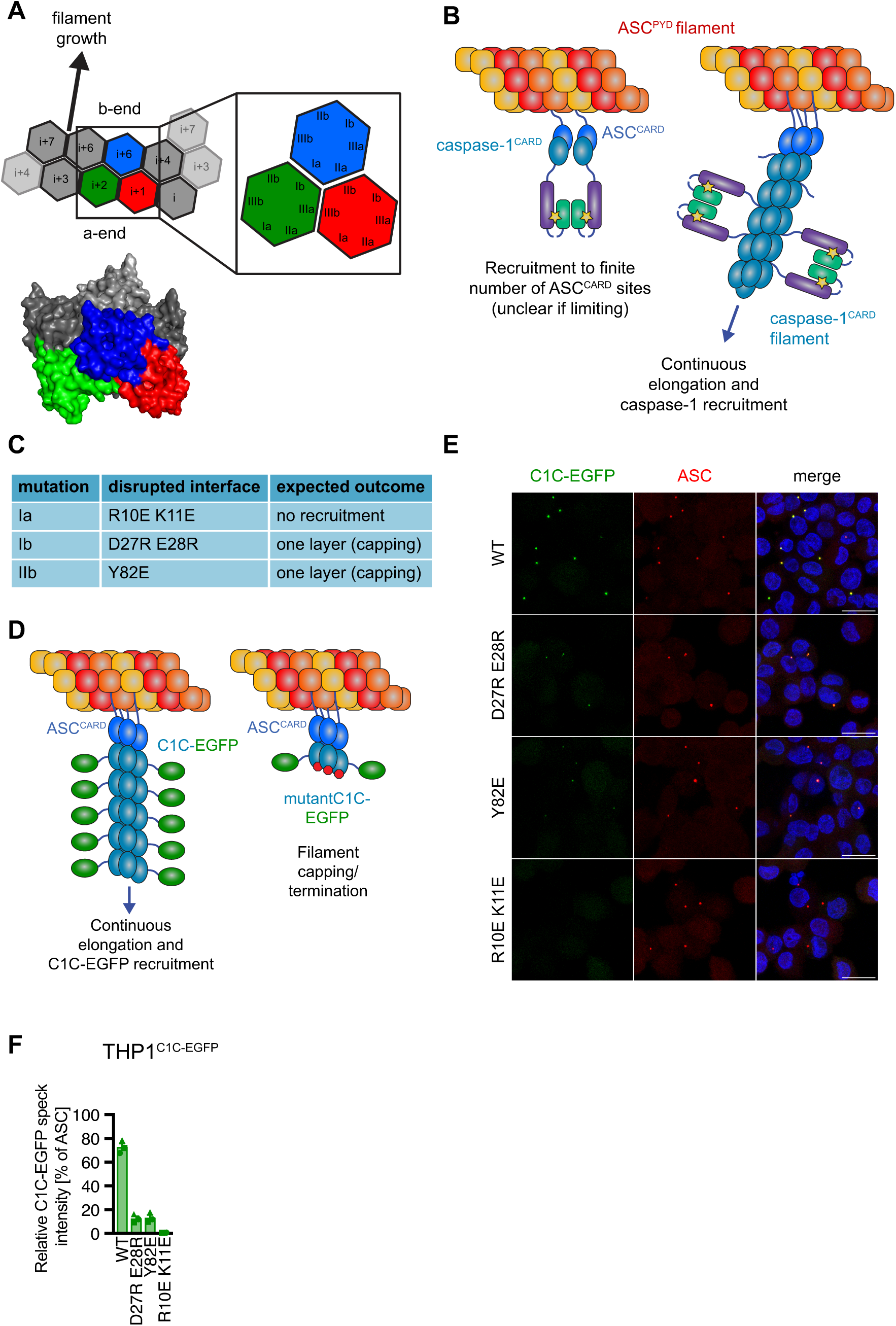
Caspase-1-EGFP is recruited to inflammasomes in cells as filaments. **A)** Schematic representation of interaction interfaces within a C1C filament as defined by Lu *et al*. (2016) (PDB 5FNA). **B)** Illustration of the models for caspase-1 recruitment as either dimers or filaments. **C)** Overview of the applied mutations to disruption C1C interaction interfaces. **D)** Illustration of the predicted termination of C1C filaments by C1C capping mutations (red octagons). **E–F)** PMA-differentiated THP-1^C1C-EGFP(i)^ or comparable cell lines inducibly expressing the indicated C1C-EGFP mutants were induced with doxycycline and treated with LPS and nigericin in the presence of VX-765 for 1 h as in Fig. 1E. Endogenous ASC was stained, and Z-stacks of cells were recorded by confocal microscopy. Representative images are displayed in (E). Scale bars: 20 μm. ASC specks were detected by Imaris based on the staining of endogenous ASC. For THP-1 cells expressing C1C-EGFP WT, D27R E28R, Y82E, or R10E K11E a total of 255, 362, 287, and 293 specks from 3 independent experiments were analyzed, respectively. The sum intensity of C1C-EGFP was measured in each speck and normalized to the respective ASC intensity (F). Data represents mean values from 3 independent experiments ± SEM.

To investigate the polymerization of caspase-1^CARD^ filaments in living cells, we generated mutants of C1C-EGFP that are defective in different interaction interfaces (Figure 5C) (Lu *et al*., 2016). As the Ia interface on C1C is necessary for the interaction with ASC^CARD^, we expect that the Ia defective mutant (C1C-EGFP R10E K11E) is not recruited to ASC specks. In contrast, we expect that C1C-EGFP mutants that are defective in their type b interfaces (Ib: D27R E28R, IIb: Y82E) are still recruited to ASC specks, because those interfaces are not involved in the interaction with ASC^CARD^, but impair recruitment of further caspase-1^CARD^ subunits in a growing filament. If caspase-1 indeed is recruited to ASC specks in the form of C1C filaments, the model predicts that type b mutants terminate (cap) those filaments and thus limit caspase-1 recruitment and processing (Figure 5D). In contrast, if individual caspase-1 molecules are merely recruited by binding to ASC^CARD^ domains in close proximity, followed by dimerization of the caspase-1 catalytic domains, the type b mutants are expected to not exert any effect.

We generated THP-1 cells expressing the different mutants of C1C-EGFP and treated cells with LPS and nigericin to activate NLRP3. We stained endogenous ASC and recorded images of the cells using confocal microscopy (Figure 5E). Using the spot detection algorithm of the image analysis software Imaris, we identified ASC specks based on the ASC staining and defined volumes of interest around each speck. We next measured the total intensity of C1C-EGFP in each volume and divided it by the respective ASC intensity to normalize the number of caspase-1 molecules that are recruited to the number of accessible ASC molecules (Figure 5F). WT C1C-EGFP was robustly recruited to ASC specks as described above and yielded a strong C1C-EGFP signal within the specks. No recruitment of the Ia mutant C1C-EGFP R10E K11E was observed, which is in line with its assumed function in the binding to ASC^CARD^. C1C-EGFP mutants with impaired b interfaces were recruited to ASC specks, but the observed intensity was substantially reduced. This perfectly fits the predictions of the model of caspase-1 recruitment that relies on C1C filaments, in which mutations in the b interfaces prevent further recruitment of monomers and thus cap the growing filaments. The b-site mutants of C1C-EGFP exhibited comparable expression levels to WT C1C-EGFP, indicating that the observed differences cannot be attributed to different expression levels (Figures S4A–B). We conclude that the strong WT C1C-EGFP signal arises from filaments with many C1C-EGP molecules nucleated by ASC^CARD^ seeds. The C1C-EGFP signal from capping mutants is diminished as only a single layer of C1C can be recruited to each ASC^CARD^ assembly. We therefore conclude that the most likely scenario is that full-length caspase-1 is recruited in a similar fashion.

### C1C filaments are a target for regulation by CARD17

The CARD-only protein (COP) CARD17, also known as INCA, is composed of a single CARD that can be recruited to C1C and caps C1C filaments *in vitro*, but cannot oligomerize (Lu *et al*., 2016). This was predicted to be caused by incompatible charges in the Ib and IIb interfaces. It can thus be described as a naturally occurring variant of C1C, which resembles the capping mutants of C1C (D27R E28R, Y82E). The model of caspase-1 recruitment that involves C1C filaments predicts CARD17 would have the capacity to terminate endogenous C1C filaments and thereby suppress caspase-1 recruitment and activation (Figure 6A).

**Figure 6:**
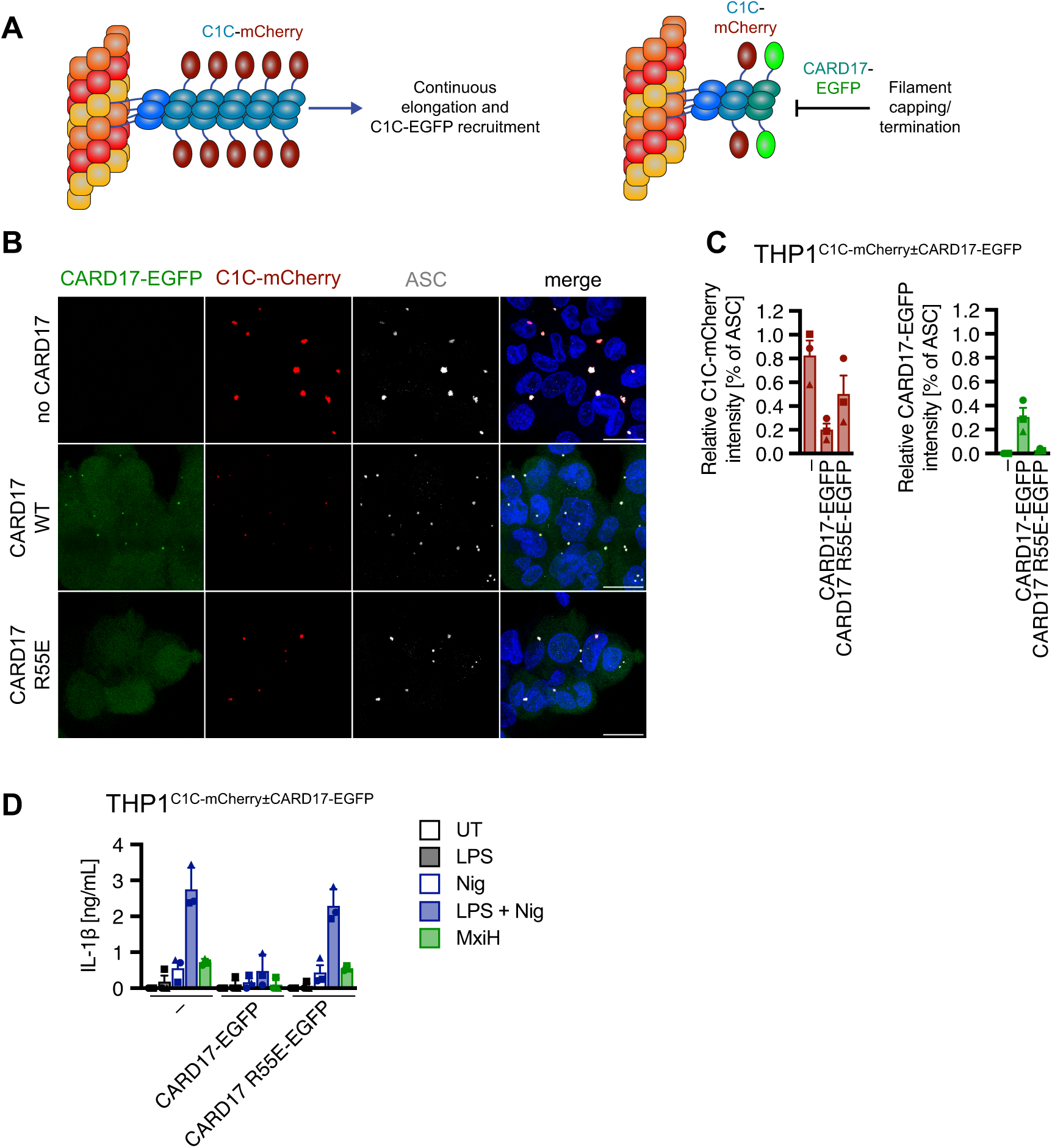
CARD17 inhibits caspase-1 by terminating filaments in cells. **A)** Illustration of the model for CARD17-mediated termination of C1C filaments. **B–C)** PMA-differentiated THP-1^C1C-mCherry^, THP-1^C1C-mCherry+CARD17-EGFP^, or THP-1^C1C-mCherry+CARD17 R55E-EGFP^ cells were treated with LPS and nigericin in the presence of VX-765 for 1 h as described in Fig. 1E. Endogenous ASC was stained, and Z-stacks were recorded by confocal microscopy. Representative maximum intensity projections are shown (B). Scale bars: 20 µm. ASC specks were identified by Imaris based on the staining of endogenous ASC. For THP-1^C1C-mCherry^, THP-1^C1C-mCherry+CARD17-EGFP^, and THP-1^C1C-mCherry+CARD17 R55E-EGFP^ a total of 1282, 1713, and 944, specks from 3 independent experiments were analyzed, respectively. The sum intensity of C1C-mCherry and CARD17-EGFP was measured in each speck and normalized to the respective ASC intensity (C). **D)** The indicated PMA-differentiated THP-1 cell lines were stimulated with LPS, LPS and nigericin, or PA + LFn-MxiH as described in Fig. 2A. IL-1β released into the supernatant was measured by HTRF. Data represents mean values from 3 independent experiments ± SEM.

To investigate the consequences of CARD17 expression to C1C recruitment, we generated derivatives of a THP-1 cell line inducibly expressing C1C-mCherry that constitutively expressed CARD17-EGFP or CARD17 R55E-EGFP, a mutant of the type Ia interface that is recruitment deficient (Lu *et al*., 2016). We then treated these cells with LPS and nigericin and recorded images by confocal microscopy (Figure 6B). As expected, C1C-mCherry was strongly recruited to ASC specks in the parental cell line lacking CARD17. In the presence of CARD17, the C1C-mCherry specks appeared much smaller. We quantified the intensity of all channels within each ASC speck as outlined above (Figure 6C). Although most of the ASC specks contained detectable amounts of C1C-mCherry, the intensity of C1C-mCherry per ASC and thus the number of recruited molecules was drastically reduced in the presence of CARD17, indicating that CARD17 indeed interfered with C1C-mCherry recruitment, likely by terminating C1C filaments (Figures 6A, S5A). CARD17-EGFP was also recruited to the specks, although the intensity was weaker, in line with the inability to oligomerize. The CARD17 mutant R55E was not recruited to ASC specks as expected, and consequently did not affect the levels of recruited C1C-mCherry. When C1C-mCherry speck formation was quantified by flow cytometry, only specks with high levels of (oligomerized) C1C-mCherry could be detected, while no C1C-mCherry specks were detected in the presence of CARD17 WT (Figure S5B). As before, mutation of the Ia interface of CARD17 restored oligomeric C1C-mCherry recruitment and speck formation.

To assess whether endogenous caspase-1 was governed by the same principles of C1C oligomerization, we next tested whether CARD17 inhibits IL-1β release, as the catalytic activity of caspase-1 required for IL-1β maturation and GSDMD pore formation is merely mediated by the endogenous protein and not the reporter C1C-EGFP (Figure 6D). Indeed, IL-1β release was nearly completely shut down by overexpressing CARD17, but not the recruitment-deficient mutant CARD17 R55E. This confirms that the same principles that govern C1C-mCherry recruitment control recruitment and activation of endogenous caspase-1. TNF release was not affected (Figure S5C), indicating that viability or priming of the cells were not affected.

In summary, we specify a mechanism for inflammasome inhibition by CARD17: effective caspase-1 recruitment and activation relies on the formation and growth of caspase-1 filaments in cells. CARD17 terminates those filaments and thus limits caspase-1 recruitment, activation, and IL-1β release. These findings also demonstrate that fluorescently labelled C1C is a powerful tool to not only recapitulate assembly of ASC specks, but also the recruitment of caspase-1.

### C1C-EGFP can indicate ASC-independent caspase-1 recruitment

In some instances, caspase-1 can be recruited and activated by inflammasome sensors in the absence of ASC. One example is CARD8, which closely resembles the active fragment of NLRP1 (UPA-CARD). Both proteins can be activated by the DPP9 inhibitor talabostat. Consequently, responses to talabostat cannot be attributed to one or the other if both are expressed. However, CARD8 recruits caspase-1 directly and cannot bind ASC (Schmidt lab, unpublished data, Ball *et al*., 2020; Gong *et al*., 2021; Hollingsworth *et al*., 2021). Human NLRP1, on the other hand, was reported not to interact with caspase-1 directly and rely on ASC to induce pyroptosis and cytokine secretion (Ball *et al*., 2020; Gong *et al*., 2021; Hollingsworth *et al*., 2021). To study the (putative) interaction of NLRP1 and CARD8 with C1C in the absence of ASC, we generated HEK 293T cells expressing either of these sensors together with C1C-EGFP, treated them with talabostat, and analyzed them by confocal microscopy (Figure 7A) and flow cytometry (Figures 7B-C). CARD8 was indeed able to nucleate the assembly of long C1C-EGFP filaments—the expected morphology if each sensor oligomer initiates a single filament rather than highly cross-linked ACS^PYD^ filaments with many potential nucleation sites for shorter C1C-EGFP filaments. To our surprise and in contrast to the previous reports, NLRP1 can also induce the assembly of C1C-EGFP in the absence of ASC, albeit less frequently. While these findings require further experiments and analysis, it is important to note that the formation of C1C-filamentous structures can also be quantified by flow cytometry using comparable gating strategies to spherical C1C specks, even though the populations of cells with and without filaments are less distinct.

**Figure 7:**
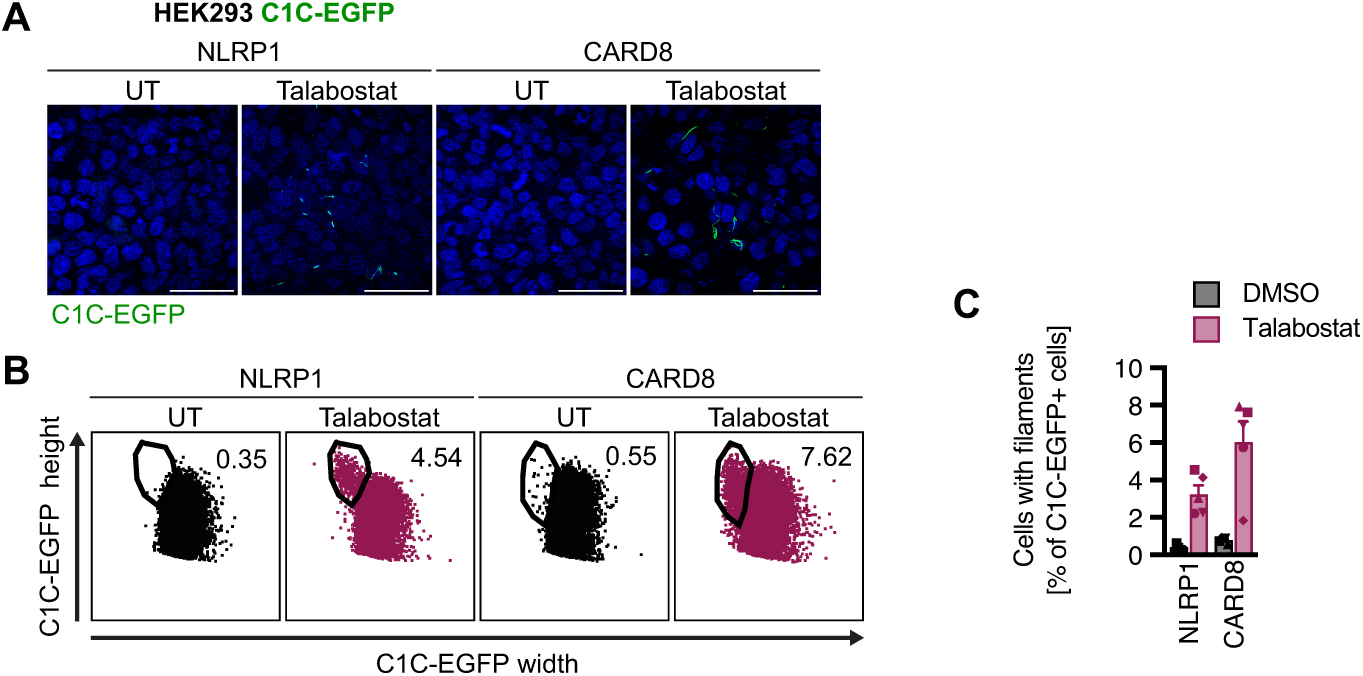
C1C-EGFP can detect filaments nucleated by CARD-containing inflammasome sensors. HEK^NLRP1-SH(i),C1C-EGFP^ or HEK^CARD8-SH(i),C1C-EGFP^ cells were treated with 30 µM talabostat for 20 h. Z-stacks of cells were recorded by confocal microscopy and representative maximum intensity projections are shown (A). Scale bars: 50 μm. The fraction of cells forming C1C-EGFP filaments was quantified by flow cytometry using a similar gating strategy as described for C1C in ASC specks in Fig. 1D. Representative dot plots (B) as well was quantified data from 5 independent experiments ± SEM is displayed (C).

These observations illustrate that C1C-EGFP also allows visualization of C1C-EGFP filaments directly recruited by CARD-containing NLRs, which will be helpful for further research on proteins like CARD8 that do not employ ASC.

## Discussion

Inflammasomes are macromolecular signaling complexes that coordinate inflammation in the tissue. Yet, most currently used tools to quantify inflammasome assembly are bulk measurements that do not reveal which and how many cells respond. We here present a novel reporter for inflammasome activation that employs a fluorescent fusion of C1C, which is recruited to assembled ASC specks. Inflammasomes can thus be localized and quantified using microscopy or flow cytometry without the need for staining. Inflammasome assembly is detected at the single cell level and can reveal responding cells in heterogeneous mixtures of different cell types. As the reporter is genetically encoded, live-cell imaging captures the dynamics of inflammasome assembly at endogenous levels of all inflammasome components. C1C-EGFP reports on the assembly of ASC specks and caspase-1 recruitment, and its recruitment thus reveals the first committed step of canonical inflammasome assembly. Importantly, incorporation of ectopically expressed C1C in inflammasomes does not affect caspase-1 activity and cytokine responses. We demonstrate that the reporter reliably detects AIM2, NLRC4, NLRP1, and NLRP3 inflammasomes in cell lines, primary cells, and organoids. We have also successfully used derivatives of the C1C reporter that use the fluorescent proteins tagBFP, mCherry, or emiRFP670 instead of EGFP (data not shown). Lastly, the reporter can be used to study the mechanistic details of caspase-1 recruitment and its regulation, as well as ASC-independent inflammasomes.

While cytokine processing and release are validated readouts for inflammasome activation, C1C-EGFP recruitment to ASC specks recapitulates both inflammasome assembly *per se* and the fraction of responding cells specifically. Using a range of nigericin concentrations in LPS-primed THP-1 cells, we observed a close correlation between IL-1β release and the fraction of specking cells, indicating that the amount of IL-1β is mostly dictated by the number of responding cells that fully commit to cytokine secretion, rather than a differential release of cytokines per cell. The secreted levels of inflammasome-related cytokines IL-1β and IL-18 also depend on the expression of sensors, cytokines and GSDMD, which can be controlled by other co-stimulatory events. This becomes apparent in nigericin-treated THP-1 cells that were not primed with the TLR4 agonist LPS. In this case, only minimal IL-1β was released despite robust inflammasome assembly. Assembly of the signaling complex is thus a more robust readout that is not affected by priming, or the expression of pro-IL-1β or pro-IL18, which differ in various tissues. Recruitment of C1C-EGFP to ASC specks is also not affected by pathogen-encoded factors that may interfere with co-stimulatory signaling pathways or any of the downstream components of the cascade. As case in point, poxviruses *e.g.* encode factors that impair transcriptional responses by NF-κB, type I interferon induction, and caspase activity, while also encoding soluble receptors for IL-1β and IL-18 (Veyer *et al*., 2017). Inflammasome assembly is thus likely the most upstream and therefore most robust and sensitive readout for inflammasome activation in infections.

To understand inflammasome biology in a physiological context, it is invaluable to study primary cells. We present a strategy to investigate inflammasome activation at the level of inflammasome assembly: primary cells that can be cultured for multiple days, including human monocyte-derived macrophages and T cells, can be transduced with lentiviruses to express C1C-EGFP prior to experiments. Unlike ectopic overexpression of ASC-EGFP applied elsewhere, C1C-EGFP expression does not alter the levels of ASC and is not prone to assembly into speck-like structures in the absence of genuine inflammasome triggers. It is thus compatible as a readout for inflammasome assembly with little background and without the need to select suitable cell clones. While this approach is limited to cells that allow transduction with lentivirus or adeno-associated viruses (AAV), it is conceivable to allow more rapid reporter expression *e.g.* by delivery of C1C-EGFP-encoding mRNAs. The need for genetic delivery of the reporter can be elegantly solved by introducing the reporter into the virus to be studied. In this way, both virus infection and inflammasome assembly can be monitored at the same time at single cell resolution and importantly can be correlated. As a proof of concept, we quantified inflammasome activation in BMDMs by the model poxvirus vaccinia virus (VACV) expressing C1C-EGFP under an early poxvirus promoter. Two manuscripts that are submitted in parallel use recombinant VACV as well as avian influenza A virus strains encoding C1C-EGFP to study inflammasome responses in human primary cells in greater detail (manuscripts attached for review). Human N/TERT-1 keratinocytes expressing C1C-EGFP have already been successfully used to unravel a novel mechanism of NLRP1 activation relevant to the human skin (L.-M. Jenster *et al*., 2023).

The next level of complexity is to investigate inflammasomes in tissues. As a model for the intestinal epithelium, we introduced mC1C-EGFP into mouse intestinal enteroids (MIEs). The reporter organoids showed that mouse IECs assemble ASC specks in response to NAIP/NLRC4 triggers as observed by fluorescence microscopy and flow cytometry. Live cell imaging confirmed that mouse IECs assemble canonical NAIP/NLRC4 inflammasome and that cell expulsion is correlated to specking cells. We also observed inflammasome assembly and cell expulsion after infection with Salmonella (Sellin *et al*., 2014; Rauch *et al*., 2017; Samperio Ventayol *et al*., 2021).

As C1C-EGFP employs the genuine recruitment domain of caspase-1, recruitment of this reporter also allowed us to address the mechanism of caspase-1 activation. Our findings with structurally defined mutants of C1C as well as CARD17, a CARD-containing negative regulator of inflammasome assembly, are consistent with a mechanism of caspase-1 recruitment that relies on the assembly of caspase-1^CARD^ filaments nucleated by ASC^CARD^ seeds. C1C oligomerization brings two unprocessed caspase-1 molecules in sufficient proximity to allow dimerization of the catalytic domains, followed by auto-proteolytic cleavage and activation. Importantly, the continuous growth of C1C filaments implies that the recruitment of additional unprocessed caspase-1 molecules can continue until all molecules are activated, as the binding sites will not be saturated. This model of activation offers an elegant way to control caspase-1 recruitment with conditionally expressed CARD-only proteins such as CARD17, which cap the growing C1C filaments (Lu *et al*., 2016). It will be exciting to investigate the physiological regulation of CARD17 as well as other regulatory mechanisms of CARD recruitment, *e.g.* by post-translational modifications of the ASC and caspase-1 CARD (Hara *et al*., 2013; Douglas and Saleh, 2020). The mechanism of activation also elegantly explains why incorporation of C1C-EGFP into endogenous caspase-1 CARD filaments does not impair the outcome of inflammasome assembly. Future derivatives of this reporter may include additional readouts for caspase-1 activity or substrate specificity.

In summary, the described C1C-EGFP inflammasome reporter will serve as a new tool to study and quantify inflammasome biology in the most physiological settings in primary cells and tissues. C1C-EGFP visualizes inflammasomes in real-time and enables readouts by microscopy and flow cytometry, *i.e.* in experimental systems that retain maximal information on the spatial organization of inflammation and the identity of the involved cell types. As demonstrated here, C1C recruitment also serves as a direct tool to dissect the regulation of caspase-1 activity by modulating CARD polymerization.

## Supporting information

Suppplementary Figures S1-S5

## Acknowledgments

We thank Zeinab Abdullah (Institute of Experimental Imunology, University of Bonn, Germany) for providing us with mCherry-expressing salmonella strains. We are grateful to Fotis Karagiannis, Vanessa Schmitt, and Christoph Wilhelm (Institute of Clinical Chemistry and Clinical Pharmacology, University of Bonn) for initial help with enteroid preparation and maintenance. We would like to acknowledge the support of Andreas Dolf and the Flow Cytometry Core Facility of the Medical Faculty, University of Bonn, for their support, services and devices funded by the Deutsche Forschungsgemeinschaft (DFG, German Research Foundation) for project number 16372401, 387335189, 387333827, and 216372545. We would also like to thank Gabor Horvath and the Microscopy Core Facility of the Medical Faculty at the University of Bonn for providing help, services and instrumentation supported by the Deutsche Forschungsgemeinschaft (DFG, German Research Foundation) for project number 388159768, and the Bundesministerium für Bildung und Forschung (BMBF, Federal Ministry of Education and Research) – ACCENT:Foerderung von Advanced Clinician Scientist im Bereich Immunopathogenese und Organdysfunktion, Gehirn und Neurodegeneration – Foerderkennzeichen: 01EO2107. We lastly thank all other members of the Schmidt lab for validating and refining the use of C1C-EGFP in inflammasome detection.

## Funding agencies

The presented work was supported by the following funding agencies: DFG Emmy Noether Programme 322568668 (F.I. Schmidt), DFG grant GRK2168-272482170 (F.I. Schmidt). DFG grant SFB1403-414786233, DFG grant TRR237-369799452 (F.I. Schmidt). DFG grant SPP1923-429513120 (F.I. Schmidt), DFG Germany’s Excellence Strategy—EXC2151– 390873048 (F.I. Schmidt), Klaus Tschira Boost Fund (KT07; F.I. Schmidt), and Independent Research Fund Denmark DFF-International Postdoc 8026-00014B (M.H. Christennsen).

