## Supplementary material for "A novel reporter to visualize and quantify endogenous inflammasomes and caspase-1 recruitment": Suppplementary Figures S1-S5

**Supplementary Figures**

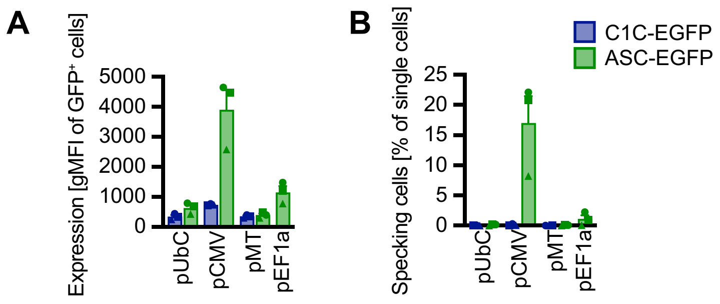


**Figure S1: C1C-EGFP allows detection of inflammasome assembly.** THP-1 cells were transduced with lentivirus encoding ASC-EGFP or C1C-EGFP under the control of the indicated constitutive promoters for 6 h and directly differentiated with PMA overnight without selection. Cells were cultivated for 24 h in the presence of 100 µM VX-765. The geometric mean fluorescence intensity (gMFI) of EGFP-positive cells (A) and the fraction of specking cells (B) was measured by flow cytometry. Data represents mean values from 3 independent experiments ± SEM.

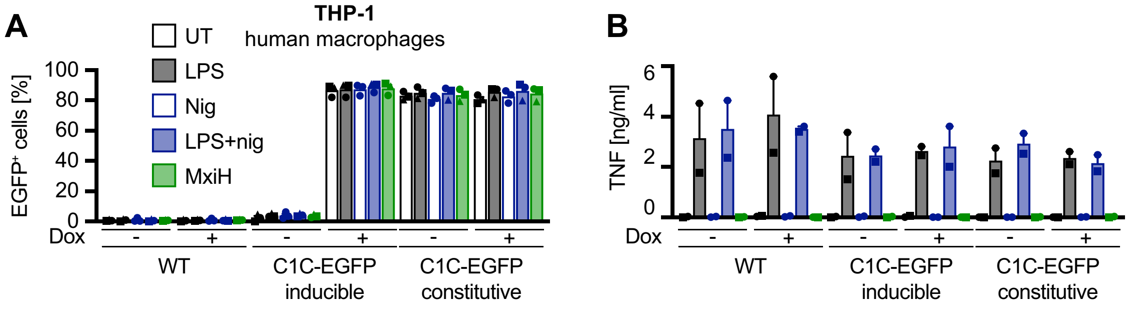


**Figure S2: C1C-EGFP does not interfere with inflammasome signaling.** PMA-differentiated WT THP-1, THP-$1^{C1C\text{-}EGFP}$, or THP-$1^{C1C\text{-}EGFP(i)}$ were treated with LPS and nigericin or with PA + LFn-MxiH for 1 h as described in Fig. 1F. The fraction of C1C-EGFP positive cells was quantified by flow cytometry (A) and TNF secretion after LPS treatment was measured by HTRF (B). The displayed data is from the same experiments that are displayed in figure 2A/B. Data represents mean values from 3 independent experiments ± SEM (2 for B).

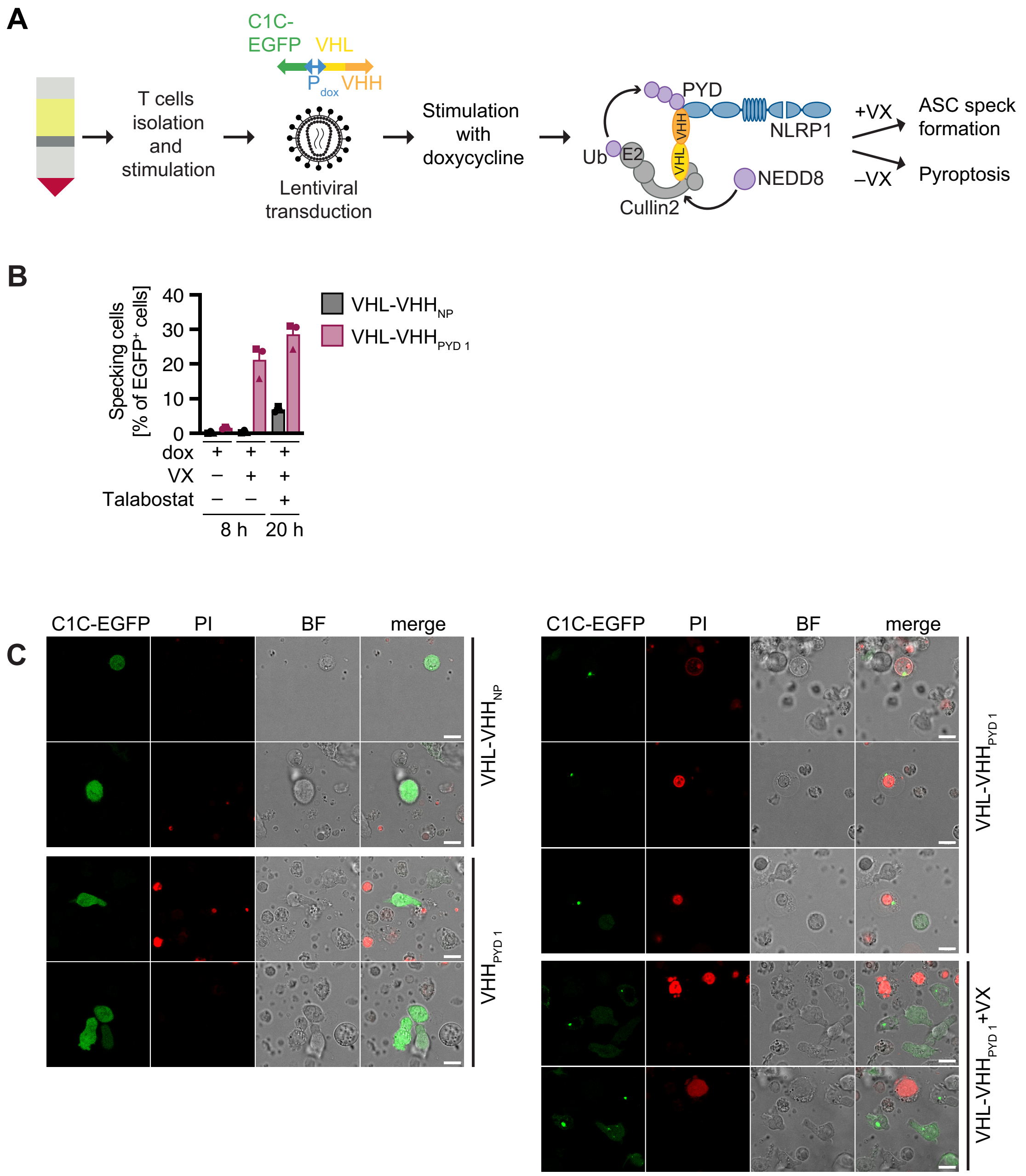

A
C1C-
EGFP
VHL
P dox
VHH
T cells
isolation
and
stimulation
Lentiviral
transduction
Stimulation
with
doxycycline
PYD
VHH
VHL
NLRP1
Ub
E2
Cullin2
NEDD8
+VX
ASC speck
formation
−VX
Pyroptosis
B
Specking cells
[% of EGFP+ cells]
40
30
20
10
0
VHL-VHH NP
VHL-VHH PYD 1
dox
VX
Talabostat
8 h
20 h
C
C1C-EGFP
PI
BF
merge
VHL-VHH NP
VHH PYD 1
VHL-VHH PYD 1
VHL-VHH PYD 1 +VX

**Figure S3: C1C-EGFP reveals inflammasome responses in primary cells. A-C)** Schematic overview of the experimental setup (A). CD3+ T cells were isolated from fresh blood, stimulated with anti-CD3/CD28 antibodies, and transduced with lentiviral vectors encoding C1C-EGFP and fusions of VHL to the indicated nanobodies (VHL-VHH), controlled by a bidirectional dox-inducible promoter. Cells were treated with 1 µg/mL dox for 8 or 20 h (B) or 6 h (C) to induce transgene expression in the presence or absence of 100 µM VX-765. Where indicated, cells were stimulated with 30 µM talabostat in the presence of VX-765 as a positive control. EGFP expression and C1C specking was quantified by flow cytometry (B). Representative images of cells induced in the presence of the non-cell permeable DNA dye propidium iodide (PI) were recorded by confocal microscopy. Scale bars represent 10 µm.

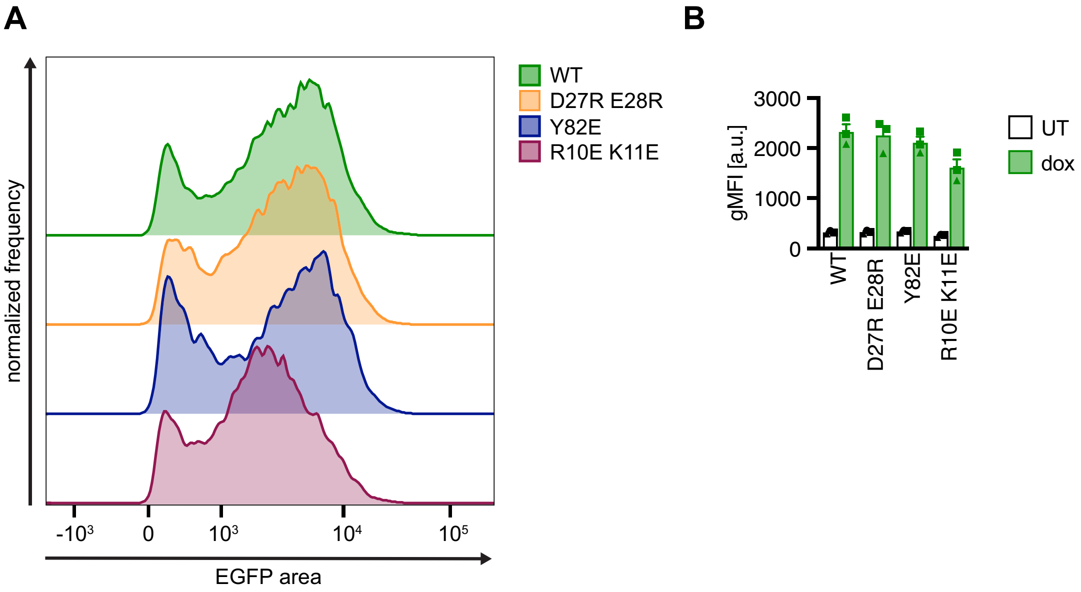


**Figure S4: Caspase-1-EGFP is recruited to inflammasomes in cells as filaments. A–B)** Undifferentiated THP-1$^{C1C\text{-}EGFP(i)}$ or comparable cell lines inducibly expressing the indicated C1C-EGFP mutants were induced with dox and C1C-EGFP expression was measured by flow cytometry. Representative histograms (A) and geometrical mean fluorescence intensity (gMFI) are displayed. Data represents mean values from 3 independent experiments ± SEM.

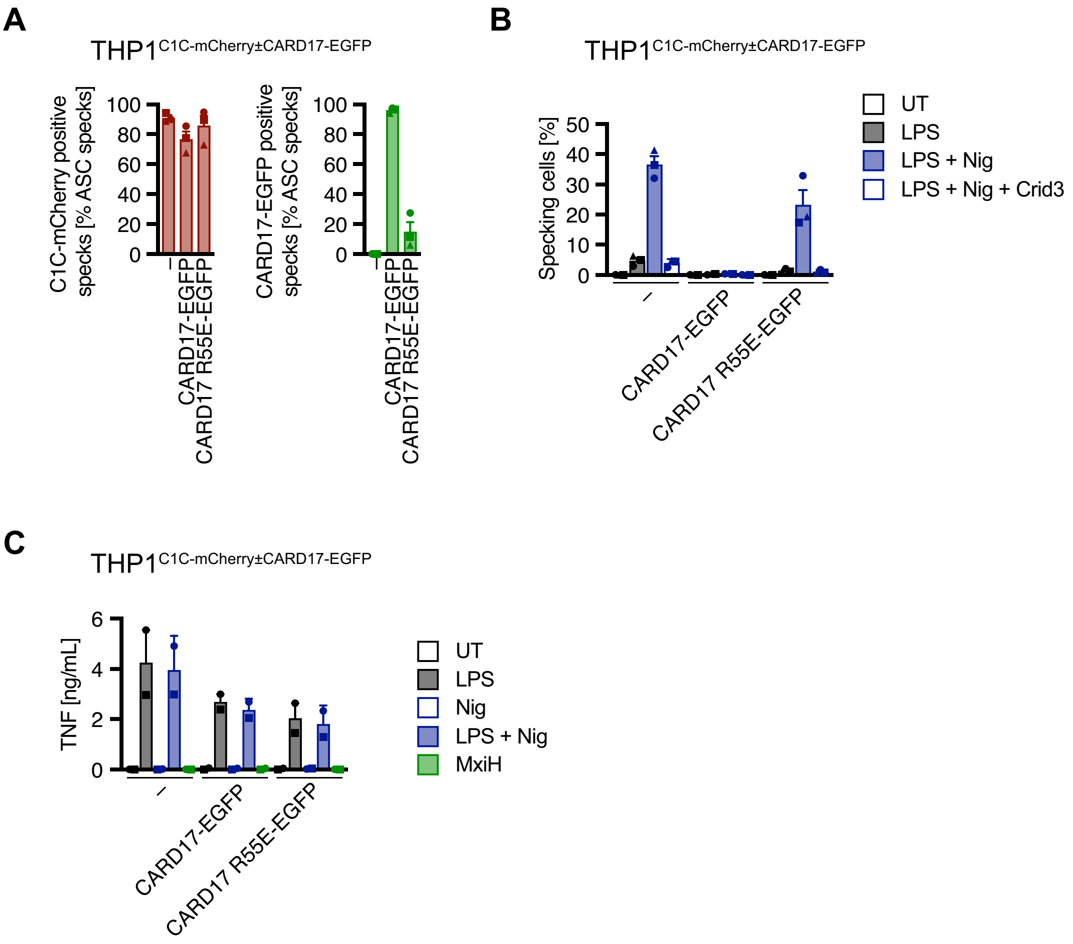


**Figure S5: CARD17 inhibits caspase-1 by terminating filaments in cells. A)** The indicated PMA-differentiatied THP-1 cell lines were induced, stimulated, and processed as described in Fig. 6B. The ASC specks detected in Fig. 6C were analyzed to determine the fraction of ASC specks that were positive for C1C-mCherry or CARD17-EGFP (sum intensity ≥ 1000). **B)** PMA-differentiated THP-1$^{\text{C1C-mCherry}}$, THP-1$^{\text{C1C-mCherry,CARD17-EGFP}}$, or THP-1$^{\text{C1C-mCherry, CARD17 R55E-EGFP}}$ cells were treated with LPS and nigericin in the presence of VX-765 for 1 h as described in Fig. 1E; where indicated cells were treated in the presence of 2.5 µM CRID3. The fraction of specking cells was measured by flow cytometry. **C)** The indicated PMA-differentiated THP-1 cell lines were stimulated with LPS, LPS and nigericin, or PA + LFn-MxiH as described in Fig. 6D. After LPS pre-treatment of samples displayed in 6D, TNF released into the supernatant was measured by HTRF. Data represents mean values from 2 independent experiments.
